# Transcription-driven cohesin repositioning rewires chromatin loops in cellular senescence

**DOI:** 10.1101/823831

**Authors:** Ioana Olan, Aled J. Parry, Stefan Schoenfelder, Masako Narita, Yoko Ito, Adelyne S.L. Chan, Guy St.C. Slater, Dóra Bihary, Masashige Bando, Katsuhiko Shirahige, Hiroshi Kimura, Shamith A. Samarajiwa, Peter Fraser, Masashi Narita

## Abstract

Senescence is a phenotypic state of stable proliferative arrest, typically occurring in lineage-committed cells and triggered by various stimuli. It is generally accompanied by activation of a secretory program (senescence-associated secretory phenotype, SASP), which modulates both local (tissue microenvironment) and systemic (ageing) homeostasis^1,2^. Enhancer-promoter interactions play a key role in gene regulation^3–5^, facilitated by chromatin loops, mostly formed via CCCTC binding factor (CTCF) and cohesin tethering^6–8^. The three-dimensional chromatin structure of senescent cells has been characterised^9–11^ mostly in terms of macro-domain structures, but its relevance in gene expression remains elusive. Here, we use Hi-C and capture Hi-C^12,13^ to show that oncogenic HRAS-induced senescence (RIS) in human diploid fibroblasts (HDFs) is accompanied by extensive enhancer-promoter rewiring, which is closely connected with dynamic cohesin binding to the genome. We find de novo cohesin peaks at the 3’ end of a subset of active genes, reminiscent of the transcription-driven ‘cohesin islands’ recently discovered in mouse embryonic fibroblasts deficient in both CTCF and the cohesin release factor Wings apart-like (Wapl)^14^. RIS de novo cohesin peaks are also transcription-dependent and enriched for SASP genes, as exemplified by *IL1B*, where de novo cohesin binding is involved in new loop formation. Cytokine induction is associated with similar cohesin islands appearance and enhancer-promoter rewiring during the terminal differentiation of monocytes to macrophages^15^, but not upon acute TNFα treatment of HDFs^16^. These results suggest that RIS represents a fate-determined process in which gene expression is regulated beyond the cell type specific 3D chromatin framework, in part through cohesin redistribution.

## Main

Oncogene-induced senescence (OIS) has been linked to dynamic alterations of the chromatin landscape, through the formation of three-dimensional heterochromatic foci^17^, altered distributions of histone modifications and chromatin accessibility^2,18^ or the appearance of new ‘super-enhancers’^19^. While it has been shown that regulation of acute stress-responsive genes can be achieved through transcription factor (TF) recruitment to largely pre-existing enhancer-promoter (EP) contacts^16,20,21^, whether or not a similar mechanism is employed during OIS (sustained oncogenic stress) is unknown. To study gene regulatory mechanisms in the 3D chromatin context at high resolution, we performed in situ Hi-C experiments as well as capture Hi-C (cHi-C) for 62 selected genomic regions of interest (Supplementary Table 1) in normal growing and oncogenic HRAS-G12V-induced senescent (RIS) IMR90 human diploid fibroblasts (HDFs), using the 4-hydroxytamoxifen (4OHT)-inducible estrogen receptor (ER) HRAS fusion system^22^ (ER:HRAS^G12V^ Fig. 1a). Growing (three replicates) and RIS (two replicates) Hi-C libraries yielded a total of 523 and 286 million valid reads respectively, after removal of artefacts and duplicates (Supplementary Table 2). There was good agreement between biological replicates, as determined with HiC-Spector^23^ and by PCA on filtered reads (cHi-C libraries were generated using DNA from the first two growing and RIS Hi-C replicates, Extended Data Fig. 1a-c).

**Figure 1:**
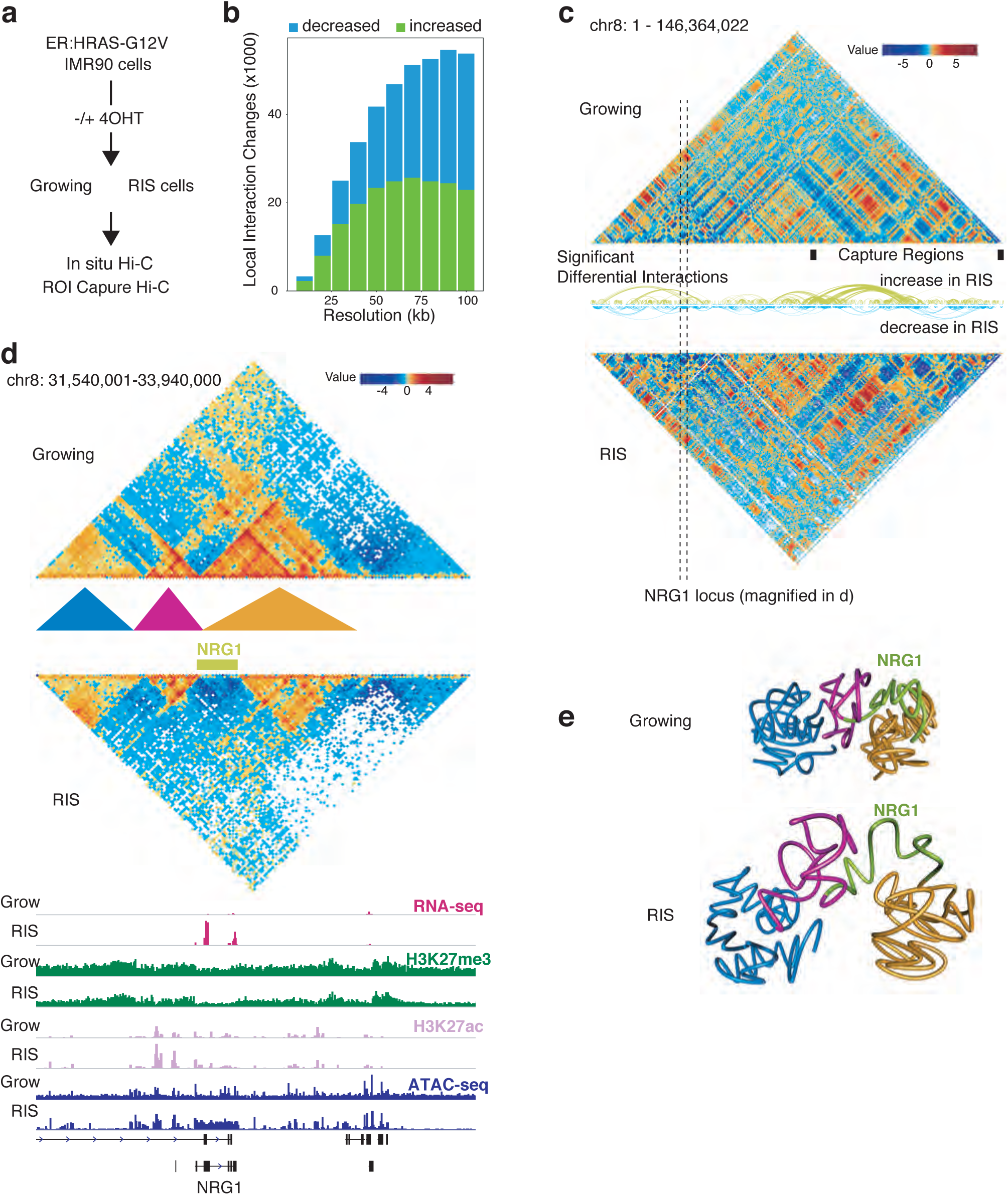
Changes in chromatin interactions identified with Hi-C during RIS: **a**, Experimental setup of control Growing and RAS-induced senescent (RIS) IMR90 cells. ROI, regions of interest. **b**, Number of significant interaction changes during RIS at resolutions between 10 and 100 kb. **c**, Hi-C matrices (300 kb resolution) of chromosome 8 in Growing and RIS cells; arcs represent significant interaction changes (100 kb resolution); black boxes represent captured regions. **d**, Hi-C interaction matrices at the *NRG1* locus, marked by dotted lines in **c** at 20 kb resolution, with matching tracks for RNA-seq, ChIP-seq, and ATAC-seq; coloured triangles represent TADs. **e**, Three-dimensional interaction modelling with TADbit at the *NRG1* locus in Growing and RIS, including *NRG1* (green) and surrounding TADs marked in **d**.

Using these Hi-C data, we identified 3,488 and 3,535 TADs in growing and RIS conditions, respectively. In agreement with a previous study^9^, TAD borders were similar between conditions (98% matched). We found virtually no differences in the distribution of A/B compartments (Extended Data Fig. 2). We then estimated differential interactions between conditions using diffHic^24^ and found extensive alterations in chromatin contacts during RIS within TADs and between distal TADs (Fig. 1b,c), similar to the previous OIS study^9^. Interestingly, the most extensive change occurred at the location of the *NRG1* gene (Fig. 1d), which was strongly up-regulated during RIS. *NRG1* was reported as a senescence marker^25^. The *NRG1* gene was largely (except for a few isoform-specific 5’ exons) encompassed in a H3K27me3-dense TAD in growing cells. However, the interactions within the gene body and with the nearby regions were almost entirely lost in RIS cells, thus releasing it from the heterochromatic TAD (Fig. 1e - TADbit modelling, see Methods). This was accompanied by an increase in chromatin accessibility across the gene, as determined by ATAC-seq (Fig. 1d). Similar behaviour was observed in the case of the *HMGA2* gene (Extended Data Fig. 1d,e), encoding a regulator of senescence-associated heterochromatic foci^26^ (SAHFs). Genome-wide, we identified 102 up-regulated genes dissociating from H3K27me3 regions in RIS (Supplementary Table 3). These data suggest that H3K27me3 regions might contribute to long-range silencing of neighbouring genes through 3D positioning within TADs and that release from such domains appears to be a relatively common mechanism of gene activation during RIS.

We next focused on gene expression and its association with regulatory elements. We annotated differential interactions with active enhancers and promoters (Extended Data Fig. 3) of genes differentially expressed during RIS^27^. We first used the high resolution (‘*HindIII* resolution’, median 4 kb) cHi-C data to identify any differential EP pairs. Within the captured regions (Supplementary Table 1), we identified 870 EP pairs that showed significantly altered interactions during RIS, involving 149 differentially expressed genes (Extended Data Fig. 4a).

To gain a genome-wide picture, we next analysed Hi-C data and identified 15,618 ‘EP interactions’, which significantly changed at 100 kb resolution. However, these EP contacts are likely to contain many false positives due to the large bin sizes compared to average enhancer or promoter size. To increase the accuracy of this estimate, we developed a strategy to filter the Hi-C EP interactions by minimising the EP changes annotated in Hi-C and not in cHi-C over captured regions (likely to be false positives), while maximising the EP changes annotated both in Hi-C and cHi-C (Extended Data Fig. 4b): enhancers with sizes greater than 7.5 kb and bin sizes smaller than 30 kb fulfilled these conditions. Using these filters, we identified 719 EP changes genome-wide from Hi-C data, involving 553 differentially expressed genes (Extended Data Fig. 5a). Combining Hi-C and cHi-C analyses, we identified 1,004 confident EP differential interactions in total (Supplementary Table 2).

The distances between interacting enhancers and promoters from both Hi-C and cHi-C were below 2 Mb, consistent with previous studies^16^. The EP network determined using cHi-C showed structures with a wide range of complexity, likely due to the high-resolution interaction information, consisting of 79 components with up to 15 nodes (enhancers or promoters). The complex rewiring was exemplified by the *IL1* and *MMP* loci, which include major SASP genes (Fig. 2a, Extended Data Fig. 4a). Although the Hi-C EP network, consisting of 479 components, was more disconnected and mostly represented a single EP interaction, the largest component consisted of 13 enhancers differentially interacting with the *INHBA* gene promoter. Of note, *INHBA* encodes a SASP factor which has been previously linked to super-enhancer activation in RIS cells^19^. Gene set enrichment analysis using genes involved differential EP interactions (Extended Data Fig. 5b) showed that transcriptionally up-regulated genes (in RIS) were significantly enriched for ‘inflammatory’ terms, whereas the down-regulated genes were enriched for ‘cell cycle’ terms. This suggests that the two senescence hallmarks, the SASP and proliferative arrest, are controlled through the rewiring of the EP network.

**Figure 2:**
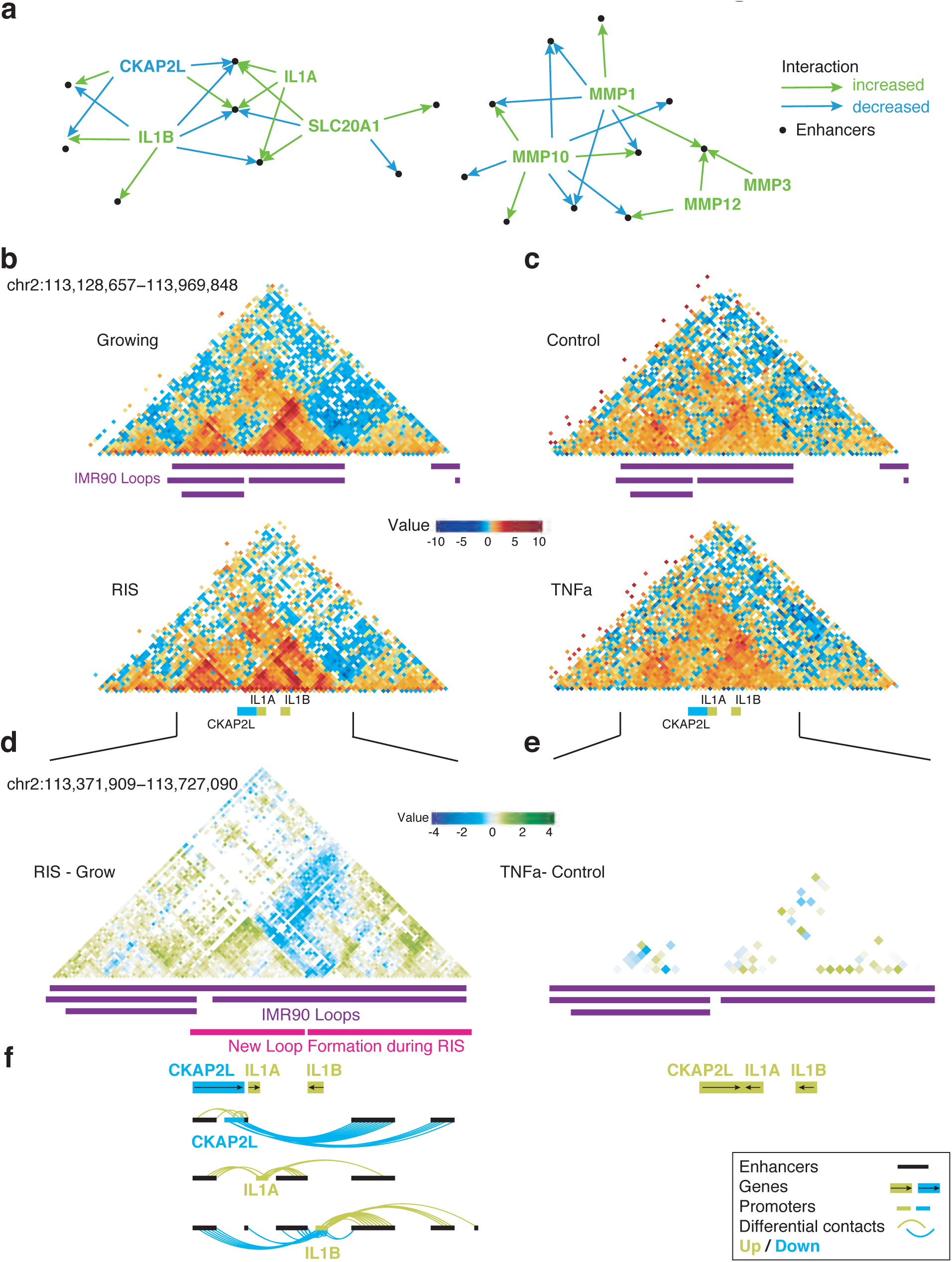
Reorganization of the local chromatin neighbourhood at the *IL1* locus: **a**, Differential enhancer-promoter contacts involving *IL1* and *MMP* in cHi-C. **b-c**, Hi-C matrices (10 kb resolution) of the *IL1* locus corresponding to growing and RIS IMR90 cells (b) and control and TNFα-treated IMR90 cells from Jin et al.^16^ (c). **d**, Differential capture Hi-C matrix at the *IL1* locus (log-fold change of RIS/growing interactions) at *HindIII* resolution, with annotated growing IMR90 loops (from Rao et al.^6^) and inferred new loop formation in RIS cells. **e**, Differential Hi-C matrix at the *IL1* locus (log-fold change TNFα) at 10 kb resolution **f**, Significant differential enhancer-promoter contacts between promoters of differentially expressed genes at the *IL1* locus and associated enhancers, aligned with d and e, respectively.

The ‘*IL1* cluster’, which was captured in our cHi-C libraries, encompasses the *IL1* ancestral family^28^ (including *IL1A*, *IL1B*) and several other genes (such as *CKAP2L*) on chr2q13. Both *IL1A* and *IL1B* encode key proximal SASP components, which are integral parts of SASP regulation^29,30^. The localization of *CKAP2L* (encoding a mitotic spindle protein) within the *IL1* cluster is highly conserved and the expression of *CKAP2L* is tightly controlled during the cell cycle^31^. Our cHi-C showed dynamic sharing of enhancers between *IL1A, IL1B*, and *CKAP2L* during RIS. The differential interaction matrix of cHi-C at the *IL1* locus showed new loop formation, segregating *IL1A* and *CKAP2L* from *IL1B* and therefore increasing the specificity of their enhancer-associations. Consistently, *IL1A* and *IL1B* began to interact more frequently with enhancers located within their respective new loops in RIS (Fig. 2f). Moreover, *CKAP2L*, which was down-regulated during RIS, interacted less frequently with the same downstream enhancers that *IL1B* began to contact more frequently (Fig. 2f). The data indicate that increased new loop formation and segregation of EP interactions occur at this locus, suggesting new loop formation around the *IL1B* gene.

This finding is in marked contrast to the *IL1* induction in a TNFα acute inflammatory scenario, in which gene regulation can be achieved without any detectable alteration to the EP landscape. Using high-resolution (5-10 kb) Hi-C maps, Jin et al.^16^ have shown that a transient TNFα treatment of IMR90 cells leads to upregulation of *IL1A* and *IL1B* with increased binding of NF-κB (a major inflammatory TF) to active enhancers of its targets. In addition, *IL1A* and *CKAP2L* were shown to be induced simultaneously via shared enhancer binding. The authors concluded that gene expression alterations mostly occur via TF binding to ‘pre-existing’ EP complexes, at least upon TNFα treatment^16^. We reanalysed the Hi-C data from this study using the presently described method and, like Jin et al., did not observe any significant changes upon TNFα treatment (Fig. 2c,e). This reveals a fundamentally distinct mechanism for the induction of inflammatory cytokines during senescence and acute inflammation. The anti-correlation between *IL1* and *CKAP2L* expression with significant EP interaction alterations was observed during RIS, but not with TNFα treatment, implying a senescence-specific decoupling mechanism within an otherwise co-regulated locus encoding key cytokines and cell cycle genes.

To investigate potential mechanisms underlying the observed EP changes during RIS, we generated ChIP-seq data for CTCF and cohesin (RAD21 and SMC3), chromatin structural proteins associated with chromatin loops^6,32^, in both growing and RIS IMR90 cells. We found 44,764 and 53,563 CTCF peaks in growing and RIS cells, respectively. Comparative analysis identified 1,774 CTCF peaks that were significantly altered during RIS. 96% of the CTCF changes were associated with increased binding in RIS (Fig. 3a). In contrast, RAD21 binding, represented by 26,374 and 24,355 peaks in growing and RIS, respectively, changed significantly at 4,553 sites, of which 81% corresponded to decreased binding (Fig. 3a). Similar results were obtained for SMC3 ChIP-seq which correlates well with RAD21 ChIP-seq signal (Fig. 3d-f, genome-wide Pearson correlation values between 0.73 and 0.96). Thus, although substantial numbers of peaks were gained in both CTCF and cohesin, a large fraction of cohesin binding was diminished.

**Figure 3:**
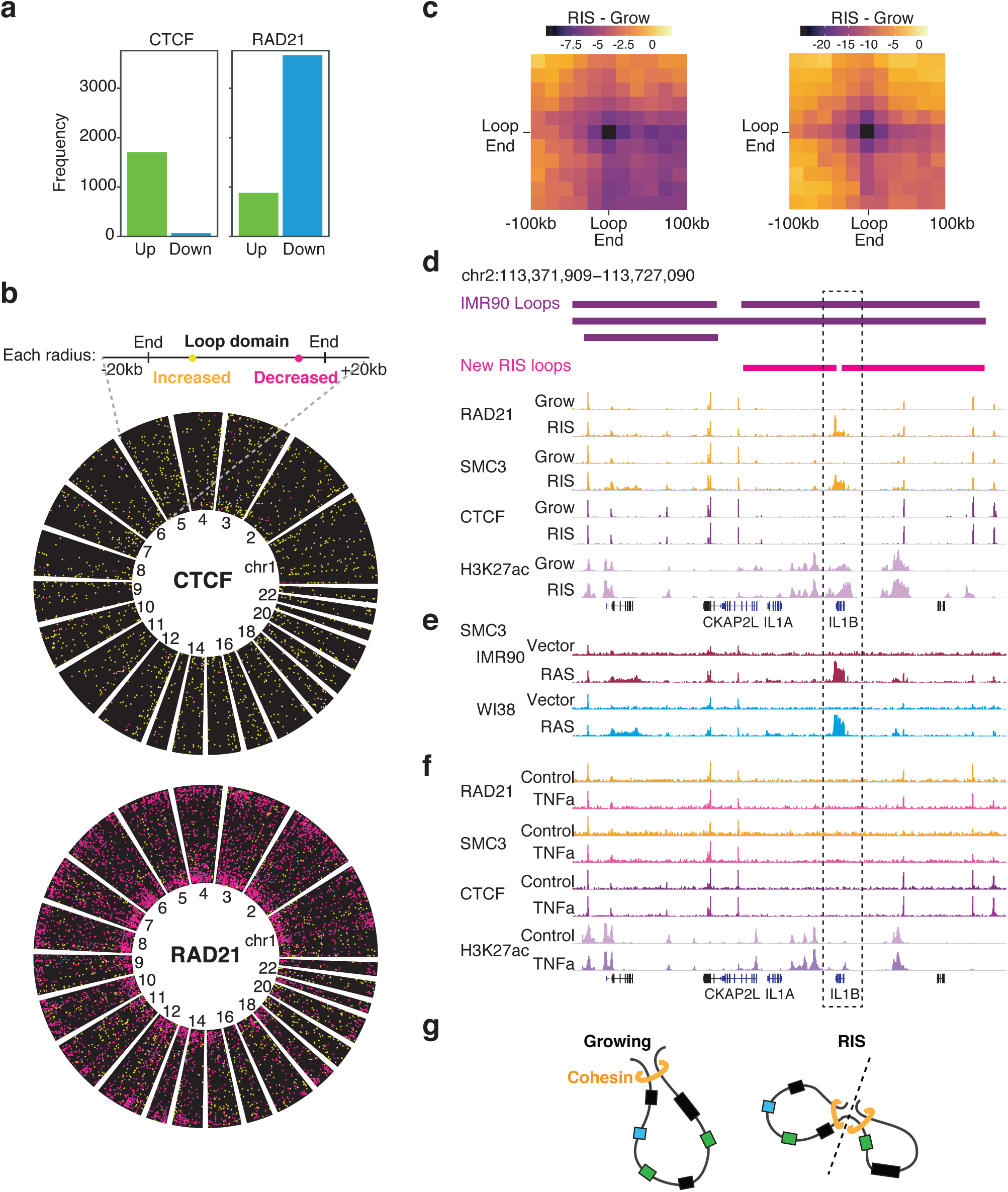
Correlation between cohesin redistribution and loop rewiring during RIS: **a**, Number of CTCF and cohesin ChIP-seq peaks with increased (green) and decreased (blue) binding. **b**, Position of CTCF and cohesin binding changes relative to the growing IMR90 loops^6^; each loop is represented as a radial segment linking the two loop anchors. **c**, Difference between RIS and growing aggregated Hi-C interactions neighbourhoods (20 kb resolution) of IMR90 loops overlapping with significantly decreased cohesin binding at least one loop ends (left). Compare to all IMR90 loops with significantly decreased interactions during RIS (right). **d**, De novo cohesin (RAD21 and SMC3) binding at the 3’ end of *IL1B* coinciding with the inferred position of the de novo loop formation in RIS, as well as CTCF and H3K27ac THOR-normalized ChIP-seq signal in growing (Grow) and RIS. **e**, THOR-normalized SMC3 ChIP-seq signal at the *IL1* locus in RIS via constitutive expression of HRAS^G12V^ (pBabe) in IMR90 and WI38 cells, as well as matched growing controls (Vector). **f**, THOR-normalized ChIP-seq signal of cohesin (RAD21 and SMC3) and CTCF at the *IL1* locus in TNFα-treated and matched control IMR90 cells. **g**, Proposed model for the de novo loop formation at the *IL1* locus, separating *IL1B* from *IL1A* and *CKAP2L*, along with their specific enhancers.

Next, we investigated where the CTCF and cohesin binding changes occurred with regards to genomic features and loops (the latter previously defined in normal IMR90 cells by Rao et al.^6^ (Fig. 3b, see Methods). 3,407 (out of 7,647) loops showed changes in cohesin binding, mostly decreases (80%), at their anchors, whereas only 363 showed any CTCF changes. Such strong colocalization between cohesin loss and loop anchors suggests that extensive loop reorganization might occur during RIS, mostly through redistribution of cohesin rather than CTCF (Fig. 3b). To visualize the relationship between cohesin reduction at loop ends and their physical contacts, we aggregated interaction neighbourhoods (at 20 kb resolution) centred on selected loops (Fig. 3c, Methods), a similar approach to the previously published method ‘Aggregate Peak Analysis’^6^. Loops with cohesin loss at one or both ends (1,827 loops) showed a trend of decreased interaction in RIS compared to growing cells (Fig. 3c, left). More stringently, 326 loops (Fig. 3c, right) were found to overlap with significantly reduced interactions during RIS, either from cHi-C or Hi-C (at 20 kb and 40 kb resolution). In terms of enhancer-promoter interactions affected, 200 differential EP contacts were nested within those robustly diminished loops, involving 92 genes, including the cell cycle regulator *CCNA2* (Extended Data Fig. 6).

Interestingly, the vast majority of cohesin binding increases occurred de novo in RIS (Extended Data Fig. 7), compared to the decreased binding which did not result in complete binding loss. The genes and enhancers studied in the *IL1* locus belong to a single loop identified in IMR90 cells^6^. This is consistent with enhancer sharing between these genes and their co-regulation in response to TNFα in these cells^16^. We found a de novo cohesin peak close to the 3’ end of *IL1B* in RIS cells, independent of CTCF binding (Fig. 3d). A similar cohesin peak was observed when RIS was induced via constitutive expression of HRAS^G12V^ without the ER-tag not only in IMR90, but also in WI38 HDFs (Fig. 3e). These data suggest that loop reorganization at the *IL1* locus is associated with de novo cohesin binding (Fig. 2d-f). Importantly, increased cohesin occupancy or altered regulatory chromatin interactions at the 3’ end of *IL1B* were not observed in response to TNFα treatment, where no new loops were detected (Fig. 2e,f, Fig. 3f). Additionally, we observed an increase in the contact intensity between the new cohesin peak and the anchors of the loop (Fig. 2d,f). Collectively, these data suggest that the de novo cohesin peak contributes to the formation of new loops in the *IL1* locus and that within each loop domain, EP pairs might preferentially contact (Fig. 3g). Strikingly, the *MMP* locus, which contains other major SASP genes, was also characterized by the appearance of de novo cohesin at the 3’ end of *MMP1* (and, to a lesser extent, *MMP3*), as well as loop reorganization around the new cohesin peak (Extended Data Fig. 8a,b). We confirmed that the cohesin increases also occurred in RIS WI38 cells (Extended Data Fig. 8c).

The elongated shape of the cohesin peaks at the 3’ end of *IL1B* and *MMP1* was reminiscent of recently reported^14^ transcription-driven ‘cohesin islands’, which appear at the 3’ end of active convergent genes in double knockout (DKO) mouse embryonic fibroblasts (MEFs) of *Ctcf* and *Wapl* (the cohesin releasing factor). The authors proposed that cohesin is loaded onto chromatin at the TSSs of a large number of active genes and is then relocated though transcription: if there is no CTCF in the way and no efficient cohesin release at the 3’ end of active genes, cohesin accumulates at the 3’ end of these genes^14^. A similar pattern of cohesin binding has been reported in wild-type yeast, which lacks a CTCF equivalent^33–35^. Thus, we hypothesized that genes highly active in RIS somehow allow for the accumulation of cohesin at their 3’ ends in a transcription-dependent manner, potentially promoting loop reorganization. To test this, we compared transcript abundance and cohesin binding at the 3’ end of genes. Both convergent (regions downstream their 3’ ends overlap) and isolated (no overlap with other genes) genes in RIS IMR90 cells accumulated cohesin islands and cohesin binding correlated with gene expression (Fig. 4a, Extended Data Fig. 9a,b). Consistent with the lack of cohesin islands detected in wild-type MEFs by Busslinger et al.^14^, very few cohesin islands were detected in normal growing IMR90 cells. To confirm that cohesin islands were associated with the cellular condition, rather than a specific subset of active genes, we examined genes highly transcribed in both RIS and growing, but at higher levels in growing cells, for cohesin islands. Despite the reduced expression levels, cohesin islands were much more pronounced in the RIS condition (Extended Data Fig. 9c).

**Figure 4.**
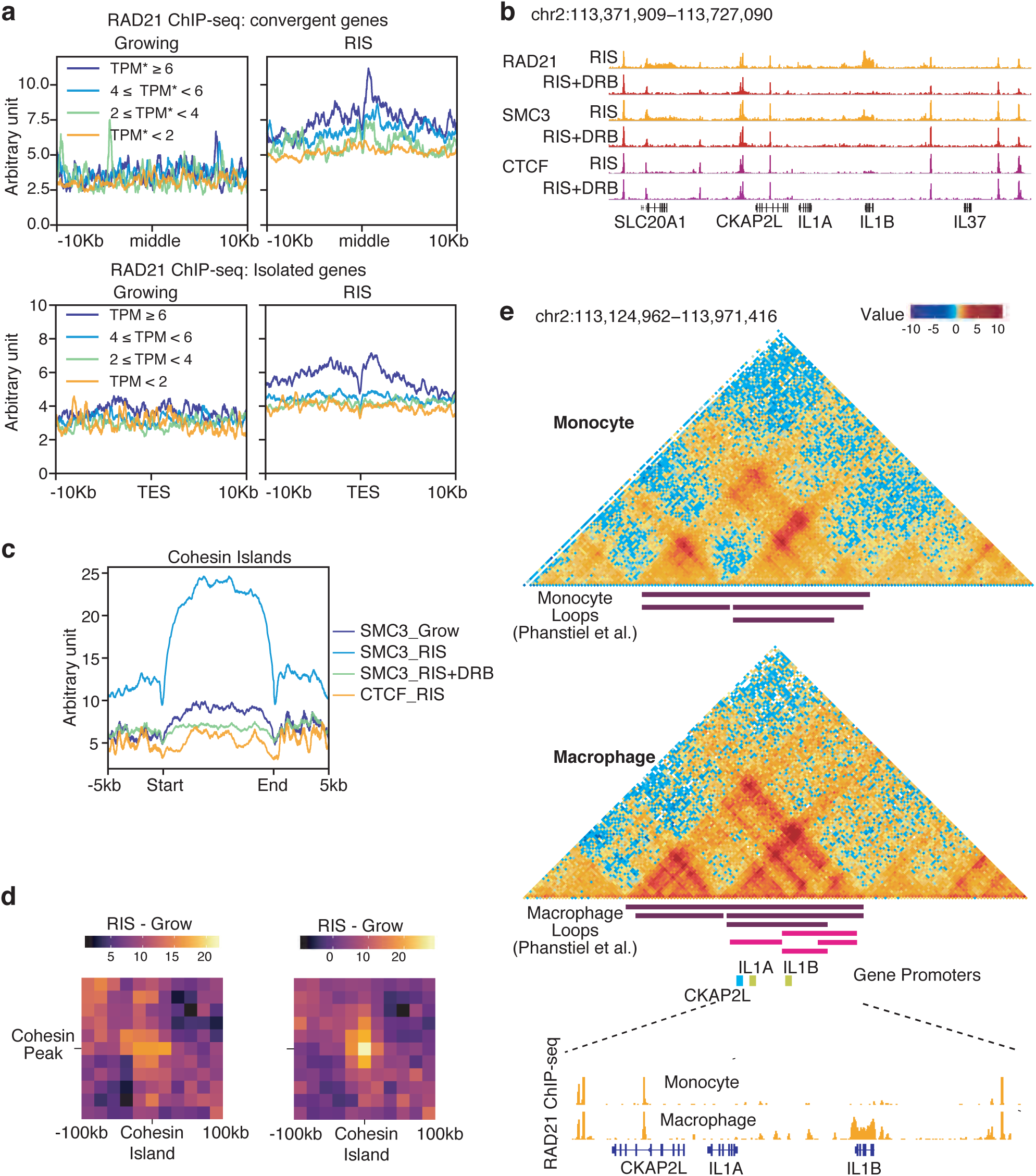
Transcription-dependent cohesin island formation in RIS associated with increased cohesin-cohesin interactions: **a**, Transcription-dependent cohesin (RAD21 THOR-normalized signal) accumulation at the 3’ end of genes in RIS cells, grouped by log-transcripts-per-million (TPM) expression at convergent genes (overlapping extended 3’ ends) and isolated genes (no overlap with other genes), with the reference points: middle point between the converging 3’ ends and TES, transcriptional end site. In the case of convergent genes, both genes in the pair were in the same expression category. **b**, THOR-normalized ChIP-seq at the *IL1* locus of cohesin (RAD21, SMC3) and CTCF with and without DRB treatment **c**, Aggregated SMC3 ChIP-seq signal in growing, RIS and RIS with DRB treatment, and CTCF ChIP-seq in RIS over all cohesin islands identified, flanked by 5 kb. **d**, Difference between aggregated interaction neighbourhoods (at 20 kb resolution) in RIS and in growing of all interactions between cohesin islands and nearby cohesin peaks within 150 kb of each other (left). The right panel represents significantly increasing interactions during RIS between each cohesin island and nearby cohesin peaks (within 250 kb either side). **e**, Cohesin islands at *IL1B* during macrophage terminal differentiation and loop modulation. Reanalysis of Hi-C matrices at 5 kb resolution of THP-1 monocytes and PMA-induced macrophages from Phanstiel et al.^15^ at the *IL1* locus, as well as cohesin ChIP-seq (RAD21) from Heinz et al.^38^ in the same cell context.

To further investigate the transcriptional dependence of cohesin islands observed in RIS cells, we performed ChIP-seq experiments in RIS cells with or without 5,6-dichloro-1-β-D-ribofuranosylbenzimidazole (DRB) treatment, a transcription elongation inhibitor. Strikingly, the de novo cohesin peaks at the *IL1B*, as well as *MMP1* sites, disappeared completely (Fig. 4b and Extended Data Fig. 8d). We used this information to define genome-wide RIS-associated cohesin islands. We found 574 wide cohesin peaks (between 2 and 20 kb wide), which were lost with DRB treatment (Fig. 4c). These regions overlapped 343 highly expressed genes in RIS, near their 3’ ends (Supplementary Table 3). In addition to *IL1B* and *MMP1* (discussed above), these genes were enriched in pathways that have been implicated in senescence, such as Wnt, Autophagy and NF-κB signalling^2,36,37^ (Extended Data Fig. 9d), and included key SASP factors. Interestingly, cohesin islands in RIS cells appeared to lack CTCF binding (Fig. 4c), suggesting that, similar to the cohesin islands defined in knockout MEFs^14^, their formation is CTCF-independent. The same cohesin islands also occurred in RIS IMR90 and WI38 cells, but not in TNFα-treated IMR90 cells (Extended Data Fig. 9e,f).

Next, we assessed whether RIS-associated cohesin islands modulate local chromatin structure, as observed at the *IL1* and *MMP* loci. Interactions between all cohesin islands and surrounding cohesin peaks (2,512 interactions) were increased in RIS (Fig. 4d, left). 105 cohesin islands exhibited significantly increased binding to local cohesin peaks (259 interactions, Fig. 4d, right). These data suggest that cohesin islands likely contribute to changes in chromatin architecture during RIS, via de novo loop formation.

Finally, we asked whether changes similar to those observed at the *IL1* locus in RIS cells occur in any other context. In most other cell types with cohesin information from ENCODE, the *IL1* locus cohesin binding pattern was similar to ‘normal’ IMR90 cells, suggesting that the loop structure at this locus is mostly conserved. However, Phanstiel et al.^15^ recently reported *IL1B* up-regulation and concomitant loop formation at the locus during terminal differentiation of monocytes into macrophages, another fate-determination process of lineage committed cells. They generated high-resolution Hi-C maps and RNA-seq datasets in the monocytic leukaemia cell line THP-1 both before (monocytes) and after (macrophages) PMA treatment, which we reanalysed. Strikingly, new loop formation in the *IL1* locus in THP-1 macrophages was similar to RIS (Fig. 4e), concomitant with similar expression changes: up-regulation of *IL1A* and *IL1B* and down-regulation of *CKAP2L*. Reanalysing RAD21 ChIP-seq data in the same THP-1 cell model from Heinz et al.^38^ revealed a de novo cohesin peak around *IL1B* (Fig. 4e). Genome-wide, cohesin binding also correlated with transcription levels (Extended Data Fig. 9g). Interestingly, 65 genes exhibited cohesin islands in both RIS and THP-1 macrophages (Extended Data Fig. 9h). Together, these data suggest that transcription-dependent cohesin accumulation also occurs during macrophage terminal differentiation and particularly, the same cohesin-mediated loop alteration at the *IL1* locus (as in RIS) facilitate transcription of genes in this locus.

We show that significantly altered EP contacts, associated with gene expression changes, occur during RIS. This is in stark contrast to proinflammatory gene expression programs in response to acute stress or signalling cues, which appear to be predominantly driven by TF recruitment and remodelling of epigenetic chromatin signatures, rather than by dynamic alteration of EP interactions. Our data indicate that EP contacts in HDFs exhibit plasticity, being susceptible to further modulation towards senescence. EP contacts in lineage-committed cells also exhibit plasticity towards terminal differentiation^15,21^. Mechanistically, our data suggest that this can be at least in part explained by the formation of transcription-dependent cohesin islands. Indeed, we also observed the induction of cohesin islands during macrophage differentiation (Fig. 4e). Generally, cohesin islands tend to be longer than the CTCF-associated structural cohesin peaks, but shorter in RIS HDFs and macrophages (up to 20 kb) than in *Ctcf*/*Wapl* DKO MEFs^14^ (up to 70 kb). Although the trigger is unclear, it is tempting to speculate that the initial de novo cohesin accumulation promotes new loop formation, and thus, increased gene expression. This would further promote transcription-dependent cohesin accumulation, constituting a gene amplification feed-forward mechanism. Our data highlight that such accumulation of cohesin islands does occur in physiological contexts in mammalian cells, where they potentially constitute an additional layer of gene regulation for cell fate determination, by modulating higher-order chromatin structure.

## Methods

### Cell culture

IMR90 and WI38 HDFs (ATCC) were cultured in Dulbecco’s modified Eagle’s medium (DMEM)/10% foetal calf serum (FCS) in a 5% O2/5% CO2 atmosphere. Cell identity was confirmed by STR (short tandem repeats) genotyping. Cells were regularly tested for mycoplasma contamination and always found to be negative. The following compounds were used in cultures: 100 nM 4-hydroxytamoxifen (4OHT) (Sigma, cat#H7904), 100 μM 5,6-dichloro-1-β-D-ribofuranosylbenzimidazole (DRB) (Sigma, cat#D1916), 10 ng/mL tumour necrosis factor alpha (TNFα) (PeproTech, cat#300-01A) as indicated in individual figures.

### Vectors

The following retroviral vectors were used: pLNCX2 (clontech) for ER:HRAS^G12V^ (Young et al.^22^), pBabe-puro for HRas^G12V^. Senescence was induced using the ER:RAS system unless otherwise mentioned.

### ChIP-seq

Chromatin immunoprecipitation (ChIP) was performed as previously described^39,40^ for the following antibodies: anti-H3K27ac^41^ (clone CMA309), anti-H3K27me3^41^ (clone CMA323), anti-CTCF (Cell Signaling Technology, clone D31H12, #3418), anti-RAD21^42^ and anti-SMC3 (Abcam ab9263). Libraries were prepared using the NEBNext Ultra II DNA Library Prep Kit for Illumina (New England Biolabs, cat#E7645L) according to the manufacturer’s instructions except that size selection was performed after PCR amplification using AMPure XP beads (Beckman Coulter, cat#A63881). Samples were sequenced paired-end using 50 bp reads on the Illumina platforms.

### Hi-C and capture Hi-C

Hi-C and capture Hi-C libraries were generated as previously described^12,13,43^ using the in-nucleus ligation protocol^44^. For each sample and replicate 50 million IMR90 cells were used. For capture Hi-C, biotinylated 120-mer RNA baits complementary to both ends of each target *HindIII* restriction fragment of interest were designed. Target sequences were valid if they contained no more than two consecutive N’s, were within 330bp of the *HindIII* restriction site and had a GC content ranging between 25 and 65%.

### Hi-C data processing

Hi-C and cHi-C libraries were aligned with HiC-Pro^45^, against the hg19 genome build. Artefacts were identified and removed using both HiC-Pro and diffHic^24^ (R Bioconductor package) and reads were counted into bins at several resolutions (*HindIII* and 5 kb for cHi-C and 10 kb - 100 kb for Hi-C). Read duplicates were removed using samtools^46^ markdup. We used HiC-Spector^23^ to check for the similarity between biological replicates, as well as PCA on library size normalized interaction matrices which were filtered for low counts and diagonal entries, produced with diffHic.

### A/B compartments

A/B compartments were called as before^47^, by performing PCA on distance-corrected, ICE-normalized Hi-C matrices at 100 kb resolution. The principal component which correlated well in absolute value with H3K4me1 ChIP-seq signal was chosen as representative of A/B compartments. The sign of the A/B compartments vector was set to match the sign of the correlation with H3K4me1 signal so that A compartment regions were represented by positive values and B compartment regions were represented by negative values.

### TADs

TADs were called using TADbit^48^ from Hi-C matrices at 40 kb resolution. A confidence score between 1 and 10 was assigned to each TAD border by TADbit. TADs from biological replicates were combined in a consensus set per condition using TADbit and only considering TAD borders with scores over 7 (out of 10).

### Interaction Modelling

We used TADbit^48^ to compute 3D models of the interactions of genes released from H3K27me3 neighbourhoods using the ICE-normalized matrices at 20 kb resolution, combined across biological replicates for growing and RIS, respectively. The matrices used correspond to the subset of interactions of one of two TADs around each gene of interest. In each case, we tried several parameter spaces for IMP parameter optimisation, employed by TADbit. For each region, we then chose the parameter subspace which fit the interaction values curve best. Modelling with IMP^49^ within TADbit was then performed with the parameters optimised for each case. The top 10 models predicted in each case were selected and exported from TADbit as XYZ coordinates.

### Differential interaction analysis

We performed differential interaction analysis between growing and RIS Hi-C and cHi-C libraries at several resolutions (*HindIII* and 5 kb for cHi-C and 10 kb - 100 kb for Hi-C, increasing in 5 kb steps) using diffHic^24^ (R Bioconductor). Libraries with artefacts and duplicates removed were further filtered for low counts and diagonal entries. Using diffHic, we performed non-linear normalization (LOESS) to remove trended biases between libraries. We tested for significant interaction changes at 5% FDR by using quasi-likelihood F-tests and Benjamini–Hochberg multiple testing correction from diffHic.

Enhancer-promoter interactions were annotated by checking the bins involved in significant differential interactions for overlaps with enhancers and promoters. We used cHi-C EP interactions annotated using *HindIII* fragments and we combined Hi-C EP interactions determined at 10 kb, 15 kb, 20 kb, 25 kb and 30 kb, filtered for enhancers longer than 7.5 kb. Enhancers were determined as before^19^, using H3K27ac peaks which overlap ATAC-seq peaks in each condition and collapsing peaks nearer than 12 kb. Promoters were represented as 5 kb regions around the TSS of protein-coding genes (GENCODE v19 reference). Only promoters of differentially expressed genes in RIS were considered.

### Using cHi-C to filter EP interactions determined with Hi-C

In order to annotate EP interactions from Hi-C more robustly, we compared several filtering strategies at different resolutions, using the contacts detected using cHi-C as a baseline for comparison, due to their accuracy at high resolution (*HindIII*). We wanted to maximise the number of EP interactions detected in the captured regions from both Hi-C and cHi-C and to minimise the interactions detected from Hi-C but not from cHi-C, which were likely false positives. We tried selecting only interactions involving enhancers of large sizes (over 5 kb, 7.5 kb or 10 kb) or genes which were more robustly differentially expressed in RIS (FDR < 0.01), as well as selecting bins without other regulatory elements. All of these filters were applied on EP interactions detected at resolutions between 10 kb and 100 kb as bin size can also affect the accuracy of the interactions detected. Finally, we selected resolutions higher than 30 kb (10 kb, 15 kb, 20 kb, 25 kb, 30 kb) and interactions involving enhancers larger than 7.5 kb for the EP interactions annotated from Hi-C.

### ChIP-seq analysis

ChIP-seq libraries were aligned against the hg19 genome build using bowtie2^50^ and uniquely mapping reads which did not bind to ‘blacklisted regions’^51,52^ were used for further analysis. Peak calling was performed using MACS2^53^ with insert sizes calculated using the R Bioconductor ChIPQC package^54^. Consensus peak sets were calculated for each condition by selecting peaks which appear in at least two replicates. Differential binding analysis was performed using THOR^55^ and a custom R script was used to filter for significant binding changes at 5% FDR and a minimum of 100 reads per location in at least one of the conditions. DeepTools^56^ was used to calculate and visualize ChIP-seq profiles summarized across genomic regions.

### RNA-seq analysis

RNA-seq libraries were aligned using STAR^57^ against the hg19 genome build and reads were counted against genes (GENCODE v19 reference) using subread^58^. Differential expression analysis was performed using glmTREAT from edgeR^59^ and significantly differentially expressed genes were selected at 5% FDR. log-TPM expression values were also calculated for the analysis of transcription and cohesin binding. Gene enrichment analysis of sets of genes of interest was performed using the enrichR R package (CRAN https://cran.r-project.org) which queries EnrichR^60,61^ against the WikiPathways database.

### Data visualization

Hi-C matrices corresponding to combined biological replicates were used for visualization. The raw matrices for each replicate were combined by calculating the overall negative binomial mean contacts with correctedContact from diffHic^24^ and further normalized using ICE^62^ and distance correction as part of the correctedContact functionality. Plotting the matrices was performed using a custom set of R scripts consisting of horizontal rotation of the matrix coordinates and overlaid genomic information such as enhancer and promoter positions, ChIP-seq tracks or interaction arcs. Interactions were coloured using a non-linear scale which represented interaction values above and below the expected values (positive and negative values, respectively, determined with distance correction) with ‘warm’ and ‘cold’ colours, respectively. In the case of overlaying cHi-C matrices at *HindIII* resolution, each interaction unit was represented as equally sized, despite the variable lengths of *HindIII* fragments. ChIP-seq tracks represented were THOR-normalized (input subtracted and library-normalized) bigWig files produced during the differential binding analysis. The tracks were exported from IGV^63^ and were scaled to be within the same interval, to allow for comparison between conditions. RNA-seq bigWig files were also produced for visualization of expression and TMM factors were used for normalization of the signal between libraries. Models derived from Hi-C interactions using TADbit^48^ were visualized using the R package rgl (CRAN https://cran.r-project.org). The points corresponding to a model are centred around 0. The curve used to visualize the model is drawn by adding 10 additional points between every pair of points in the set of original coordinates by interpolation with the spline function in R.

### Cohesin Islands

We investigated the association between cohesin accumulation and transcription by grouping genes by expression level (represented as log-TPM averaged across biological replicates) and plotting their SMC3 and RAD21 ChIP-seq profile. The binding profile was centred either at their transcription end site (TES) in the case of isolated genes or in the middle of the genomic region bounded by TES of two convergent genes. We focused this analysis on isolated and convergent genes, like in Busslinger et al.^14^, in order to avoid biases caused by genes with overlapping regions post 3’UTR. Small increases in signal were observed before the TES as well caused by short genes which show cohesin binding on their gene bodies as well.

We determined cohesin islands by comparing the RIS cohesin ChIP-seq libraries with and without DRB treatment (transcription elongation inhibitor). We then selected significantly differentially bound regions larger than 2 kb. We eliminated possible false positives which can occur due to genes with overlapping 3’ end regions (such as convergent genes) by filtering for highly expressed genes. Genes with cohesin islands were determined by overlapping 10 kb regions starting at the TES with the cohesin islands determined.

### Interaction neighbourhood aggregation

We identified general trends of certain subsets of interactions by selecting a two-dimensional neighbourhood around each interaction of interest, and summing the corresponding Hi-C sub-matrix (from the ICE-normalized and distance-corrected matrix averaged across replicates, as described earlier), similarly to Aggregate Peak Analysis^6^. Each interaction pixel was divided by the number of sub-matrices added minus the number of missing values. We selected a 200 kb region around each bin containing a cohesin peak of interest which, at 20 kb resolution, resulted in neighbourhoods of 11×11 pixels. Differential aggregated matrices were computed by subtracting the growing-specific aggregated matrix from the RIS one.

### Data availability

Hi-C and cHi-C data in growing and senescent IMR90 cells, as well as ChIP-seq data in IMR90 and WI38 human diploid fibroblasts in the growing (with and without TNFα treatment) and RIS (with and without DRB treatment) conditions were deposited in the Gene Expression Omnibus: GSExxxxx (will be made public upon publication). Publicly available ChIP-seq data were also reanalysed in this study: RAD21 and CTCF ChIP-seq in monocyte (THP-1) and macrophage (PMA-induced) controls from Heinz et al.^38^ (GSE103477), H3K4me3 and H3K27me3 ChIP-seq from Chandra et al.^64^ (GSE38448), H3K27ac ChIP-seq and ATAC-seq from Parry et al.^65^ (GSE103590). Publicly available RNA-seq data from Hoare et al.^27^ (GSE72404), Phanstiel et al.^15^ (GSE96800) and Jin et al.^16^ (GSE43070) were reanalysed in this study. Publicly available Hi-C data from Phanstiel et al.^15^ (PRJNA385337) and Chandra et al.^9^ (PRJEB8073) were also reanalysed in this study.

### Code availability

Custom scripts used for enhancer-promoter annotation and filtering THOR differential binding output were uploaded to the OSF public repository (will be made public upon publication). Visualisation scripts for the Hi-C matrices are also available on GitLab (https://gitlab.com/ilyco/hicvizr).

## Acknowledgements

We thank all members of the Narita laboratory for helpful discussions, CRUK-CI core facilities (Genomics, Biorepository, Bioinformatics and Research Instrumentation) for technical support. This work was supported by a Cancer Research UK Cambridge Institute Core Grant (C9545/A29580) to the M.N. laboratory. M.N. was also supported by the Medical Research Council (MR/M013049/1) and BBSRC (BB/S013466/1). I.O. was supported by Wellcome Trust (105367/Z/14/Z). S.S. was supported by the Biotechnology and Biological Science Research Council UK (BB/J004480/1), a Career Progression Fellowship from the Babraham Institute, and an MRC UKRI Rutherford Fund Fellowship. A.J.P. was supported by a Sir Henry Wellcome Postdoctoral Fellowship (215912/Z/19/Z). H.K. was supported by JSPS KAKENHI (JP17H01417 and JP18H05527) and JST-CREST (JPMJCR16G1). K.S. and M.B. were supported by Grant-in-Aid for Scientific Research on Innovative Areas (15H05976). S.A.S and D.B. are supported by the Medical Research Council (MC_UU_12022/10).

## Author contributions

I.O. and M.N. conceived the study. A.J.P., S.S. and YI performed Hi-C and capture Hi-C experiments. P.F. supervised Hi-C/capture Hi-C experiments and interpreted the data. Masako N. and A.J.P. performed the ChIP-seq experiments with help by H.K., K.S., and M.B. G.St.C.S. performed three-dimensional interaction modelling and visualization. I.O. analysed the Hi-C, cHi-C, ChIP-seq and RNA-seq data with input from D.B., S.A.S., and A.S.L.C. M.N. and I.O. wrote the manuscript with input from the other authors.

---

**Extended Data Figure 1. Consistency across Hi-C and cHi-C biological replicates: a**, HiC-spector agreement scores between each pair of growing and RIS biological replicates between 0 and 1, indicating poor and good agreement, respectively. **b-c**, Principal Component Analysis on read counts, which were library size-normalized and filtered for low values, corresponding to interactions at 40 kb resolution from the growing and RIS Hi-C (b) and cHi-C (c) libraries. **d**, Growing and RIS interactions matrix at 20 kb resolution of the *HMGA2* gene and surrounding TADs with matching RNA-seq (TMM-normalized) and ChIP-seq tracks of H3K27me3 and H3K27ac, as well as ATAC-seq. **e**, TADbit three-dimensional modelling of the three TADs marked in **d**, colouring the *HMGA2*, *IRAK3* and *GRIP1* genes as well as the TADs they belong to.

**Extended Data Figure 2. A/B compartments in growing and RIS:** Distribution across each chromosome of the principal component corresponding to A/B compartments (based on correlation with H3K4me1 signal) from PCA performed on growing (G) and RIS Hi-C libraries; positive values (green) mark the A compartment, whereas negative values (blue) mark the B compartment.

**Extended Data Figure 3. Epigenetic characterization of enhancers and promoters using ATAC-seq and H3K27ac, H3K4me1 and H3K4me3 ChIP-seq signal: a**, Enhancers defined by H3K27ac peaks which also have ATAC-seq and H3K4me1 and low H3K4me3 ChIP-seq signal, split by regions common between growing (G) and RIS and specific to each condition. **b**, Promoters defined as 5 kb regions around the TSS of every protein-coding gene, showing low H3K4me1 and high H3K4me3 signal.

**Extended Data Figure 4. Enhancer-Promoter (EP) Network from capture Hi-C: a**, Differential EP interactions network based on annotated cHi-C significant interaction changes at *HindIII* resolution. Two boxed components are also shown in Fig. 2a. **b**, comparison between EP interactions annotated from genome-wide Hi-C analysis at resolutions between 10 kb and 100 kb, and EP interactions annotated from cHi-C (represented in **a**) filtered with different strategies such as larger enhancers or FDR threshold of the gene; left - percentage of EP annotated from Hi-C but not from cHi-C (‘false positives’) in the captured regions, and right - percentage of EP annotated from Hi-C as well as from cHi-C (‘true positives’); the filtering strategy highlighted (enhancers larger than 7.5 kb) and bin sizes smaller than 30 kb minimise the EP annotated from Hi-C but not from cHi-C and maximise the ones annotated.

**Extended Data Figure 5. Genome-wide Hi-C EP differential network: a**, EP network using the filtering strategy based on EP changes annotated from cHi-C (Extended Data Fig. 4b, enhancers larger than 7.5 kb and resolution higher than 30 kb, i.e. bin sizes smaller than 30 kb) on each chromosome, with the vertical axis representing enhancers and the right and the left axes corresponding to up-regulated and down-regulated genes, respectively. **b**, Gene enrichment with EnrichR against the WikiPathways 2019 database of up-regulated (left) and down-regulated (right) genes in the genome-wide Hi-C EP differential network; the dotted grey line corresponds to 0.05 adjusted p-value, the selected threshold for significant enrichment. Terms were manually simplified and obvious redundancy was removed.

**Extended Data Figure 6. Down-regulation of *CCNA2* associated with decreased EP interactions:** Growing and RIS Hi-C matrices (10 kb resolution) centred on the IMR90 loop^6^ consisting of the *CCNA2* gene promoter and associated enhancers, as well as cHi-C differential log-fold change matrix (5 kb resolution) of this loop and significant decreased interactions (blue arcs) between the *CCNA2* gene promoter and associated enhancers.

**Extended Data Figure 7. RAD21 binding gains and losses in growing and RIS:** Profiles and heatmaps of THOR-normalized RAD21 ChIP-seq signal in growing and RIS, centred on peaks overlapping regions with significantly increased or decreased binding, as determined with THOR at FDR 0.05.

**Extended Data Figure 8: Interaction changes during RIS at the *MMP* locus suggesting increased spatial separation around *MMP1*: a**, cHi-C differential interaction matrix (*HindIII* resolution) at the *MMP* locus, consisting of the promoters of *MMP10*, *MMP1*, *MMP3*, *MMP12* and associated enhancers, as well as the significant EP contacts between them (green and blue arcs, corresponding to increased and decreased interactions, respectively). **b**, ChIP-seq THOR-normalized tracks at the *MMP* locus marked in **a**, with dotted lines of RAD21, SMC3, CTCF and H3K27ac in growing and RIS, as well as the *MMP* genes. **c**, SMC3 CHIP-seq of the *MMP* locus in RIS IMR90 and WI38 cells via constitutive expression of oncogenic HRAS-G12V and matched growing controls (empty vector). **d**, RAD21, SMC3 and CTCF ChIP-seq in RIS with and without DRB treatment (transcription elongation inhibitor) at the *MMP* locus with cohesin islands at the end of *MMP1* and *MMP3*.

**Extended Data Figure 9: Cohesin islands in RIS IMR90 cells and THP-1 macrophages: a**, Distribution of SMC3 ChIP-seq signal between the 3’ ends of convergent gene pairs, where both genes are in one of the four expression categories defined based on their log-transcripts-per-million (TPM* signifies TPM of both genes in the pair) in growing and RIS, respectively. **b**, SMC3 signal around 3’ end (TES) of isolated genes averaged by four expression groups defined by log-TPM of the genes in growing and RIS, respectively. **c**, RAD21 and SMC3 ChIP-seq signal at the 3’ end of 559 genes which are highly expressed in growing and RIS but more highly in growing. **d**, Gene enrichment pathways of genes with cohesin islands against WikiPathways 2019 (dotted line corresponds to the significance threshold, 0.05 p-adjusted). **e**, SMC3 ChIP-seq signal of constitutive HRAS-G12V-induced senescence and matched controls (empty vector), in IMR90 and WI38 cells, over the cohesin islands identified from RIS compared with RIS with DRB treatment. **f**, SMC3 ChIP-seq signal of RIS, TNFα-treated and matched control IMR90 cells over the cohesin islands. **g**, RAD21 ChIP-seq by Heinz et al.^38^ around the 3’ end of genes grouped by expression levels (RNA-seq from Phanstiel et al.^15^) in THP-1 monocytes and PMA-induced macrophages. **h**, Monocyte and macrophage RAD21 ChIP-seq by Heinz et al.^38^ over the cohesin islands defined in RIS IMR90 cells.

**Extended Data Table 1:** List of captured regions with hg19 coordinates

**Extended Data Table 2:** Number of valid reads aligned from each Hi-C and cHi-C library, as well as the number of reads corresponding to various types of artefacts

**Extended Data Table 3:** Genes up-regulated during RIS which also show reduced interactions with H3K27me3 regions, escaping heterochromatic three-dimensional neighbourhoods

**Extended Data Table 4:** Enhancer-promoter differential contacts from cHi-C (*HindIII* resolution) and Hi-C (at different resolutions)

**Extended Data Table 5:** Genes with cohesin at their 3’ends during RIS

