## Supplementary Figures for "Transcription-driven cohesin repositioning rewires chromatin loops in cellular senescence"

### Extended Data Fig. 1_Olan et al.

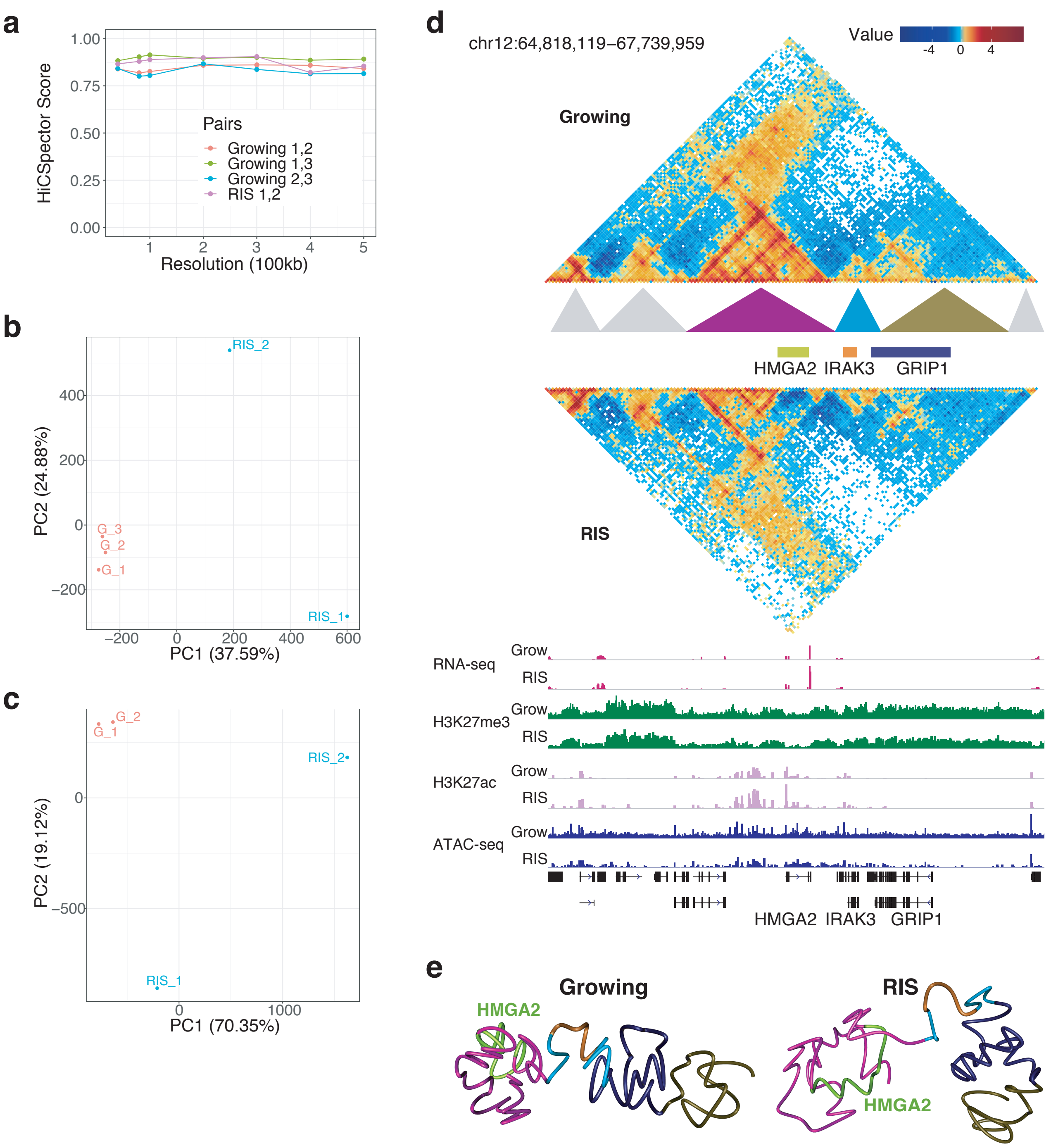


**Extended Data Figure 1. Consistency across Hi-C and cHi-C biological replicates: a,** HiC-spector agreement scores between each pair of growing and RIS biological replicates between 0 and 1, indicating poor and good agreement, respectively. **b-c**, Principal Component Analysis on read counts, which were library size-normalized and filtered for low values, corresponding to interactions at 40 kb resolution from the growing and RIS Hi-C (**b**) and cHi-C (**c**) libraries. **d**, Growing and RIS interactions matrix at 20 kb resolution of the *HMGA2* gene and surrounding TADs with matching RNA-seq (TMM-normalized) and ChIP-seq tracks of H3K27me3 and H3K27ac, as well as ATAC-seq. e, TADbit three-dimensional modelling of the three TADs marked in **d**, colouring the *HMGA2*, *IRAK3* and *GRIP1* genes as well as the TADs they belong to.

#### Extended Data Fig. 2_Olan et al.

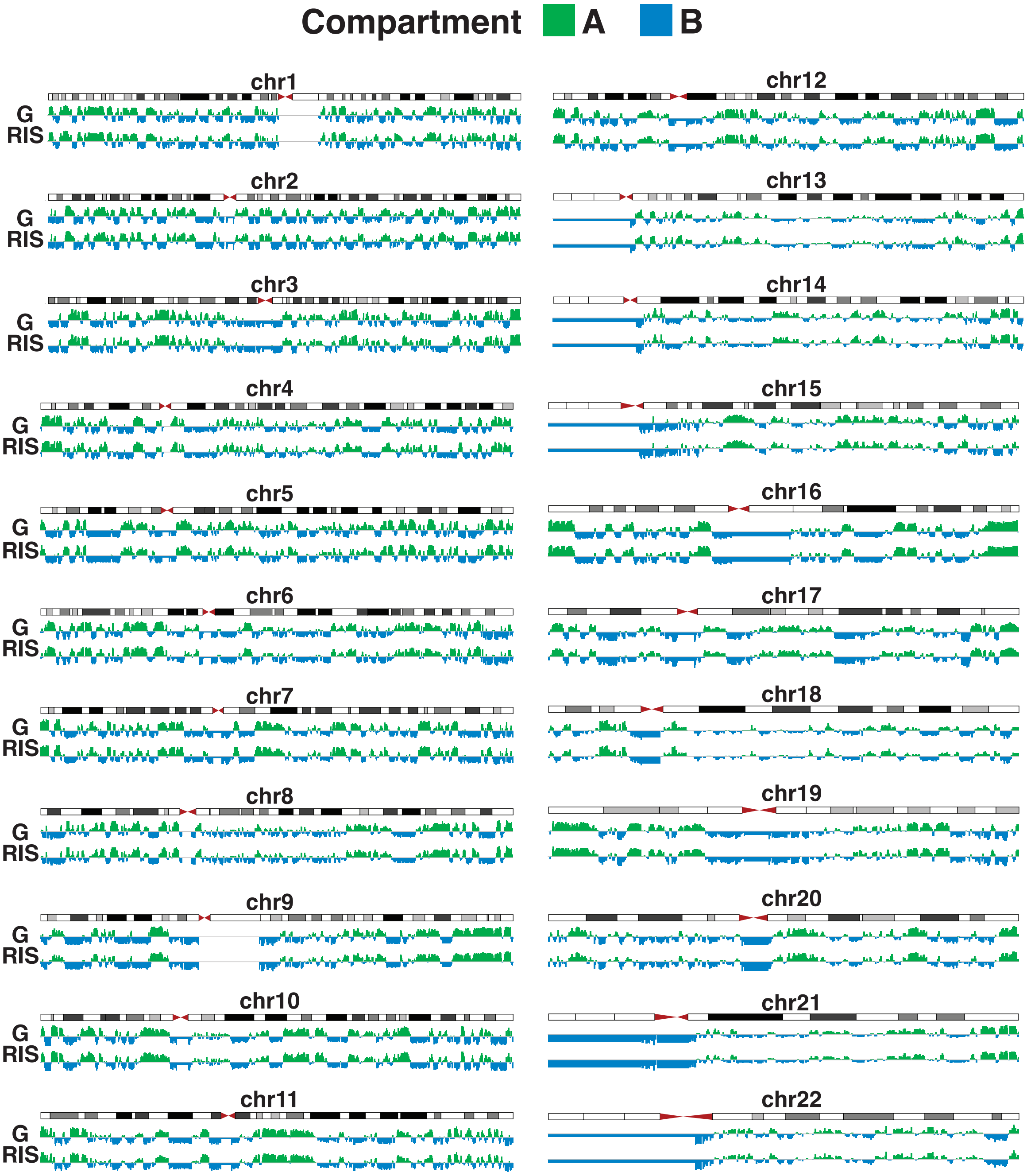


**Extended Data Figure 2. A/B compartments in growing and RIS**: Distribution across each chromosome of the principal component corresponding to A/B compartments (based on correlation with H3K4me1 signal) from PCA performed on growing (G) and RIS Hi-C libraries; positive values (green) mark the A compartment, whereas negative values (blue) mark the B compartment.

### Extended Data Fig. 3_Olan et al.

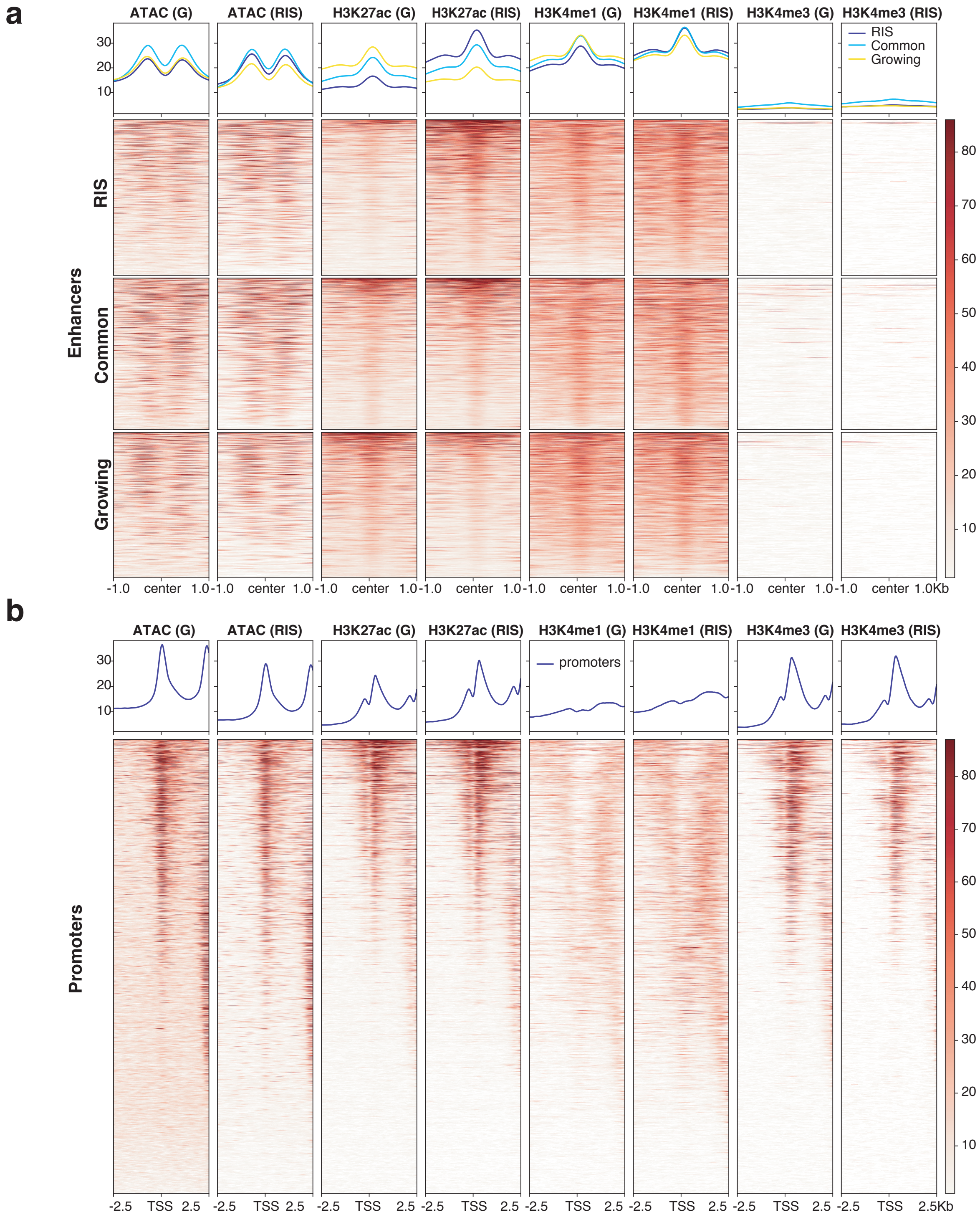


**Extended Data Figure 3. Epigenetic characterization of enhancers and promoters using ATAC-seq and H3K27ac, H3K4me1 and H3K4me3 ChIP-seq signal: a,** Enhancers defined by H3K27ac peaks which also have ATAC-seq and H3K4me1 and low H3K4me3 ChIP-seq signal (Parry et al.), split by regions common between growing (G) and RIS and specific to each condition. **b**, Promoters defined as 5 kb regions around the TSS of every protein-coding gene, showing low H3K4me1 and high H3K4me3 signal.

### Extended Data Fig. 4_Olan et al.

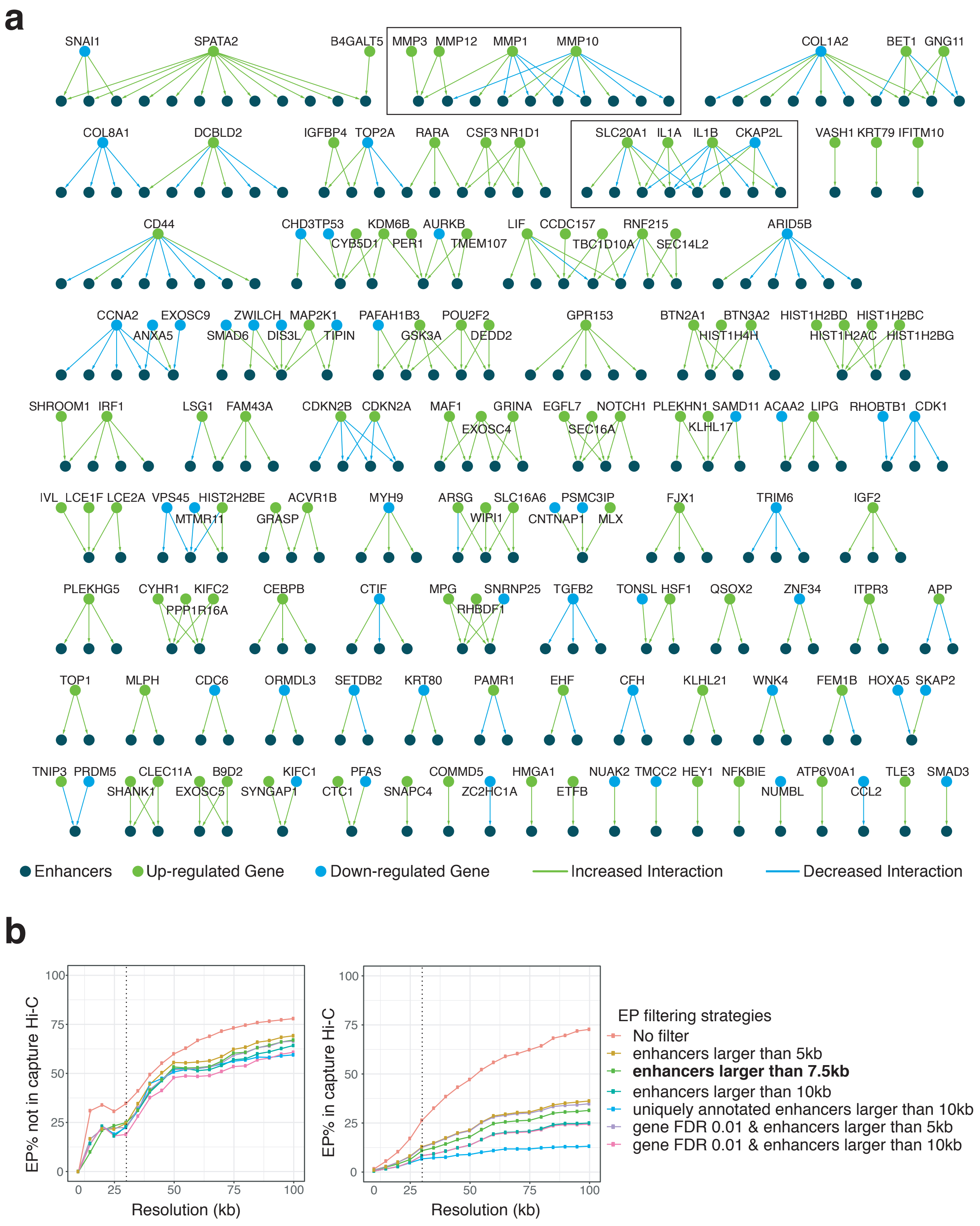


**Extended Data Figure 4. Enhancer-Promoter (EP) Network from capture Hi-C: a**, Differential EP interactions network based on annotated cHi-C significant interaction changes at HindIII resolution. Two boxed components are also shown in Fig. 2a. **b**, comparison between EP interactions annotated from genome-wide Hi-C analysis at resolutions between 10 kb and 100 kb, and EP interactions annotated from cHi-C (represented in **a**) filtered with different strategies such as larger enhancers or FDR threshold of the gene; left - percentage of EP annotated from Hi-C but not from cHi-C ('false positives') in the captured regions, and right - percentage of EP annotated from Hi-C as well as from cHi-C ('true positives'); the filtering strategy highlighted (enhancers larger than 7.5 kb) and bin sizes smaller than 30 kb minimise the EP annotated from Hi-C but not from cHi-C and maximise the ones annotated in both datasets.

### Extended Data Fig. 5_Olan et al.

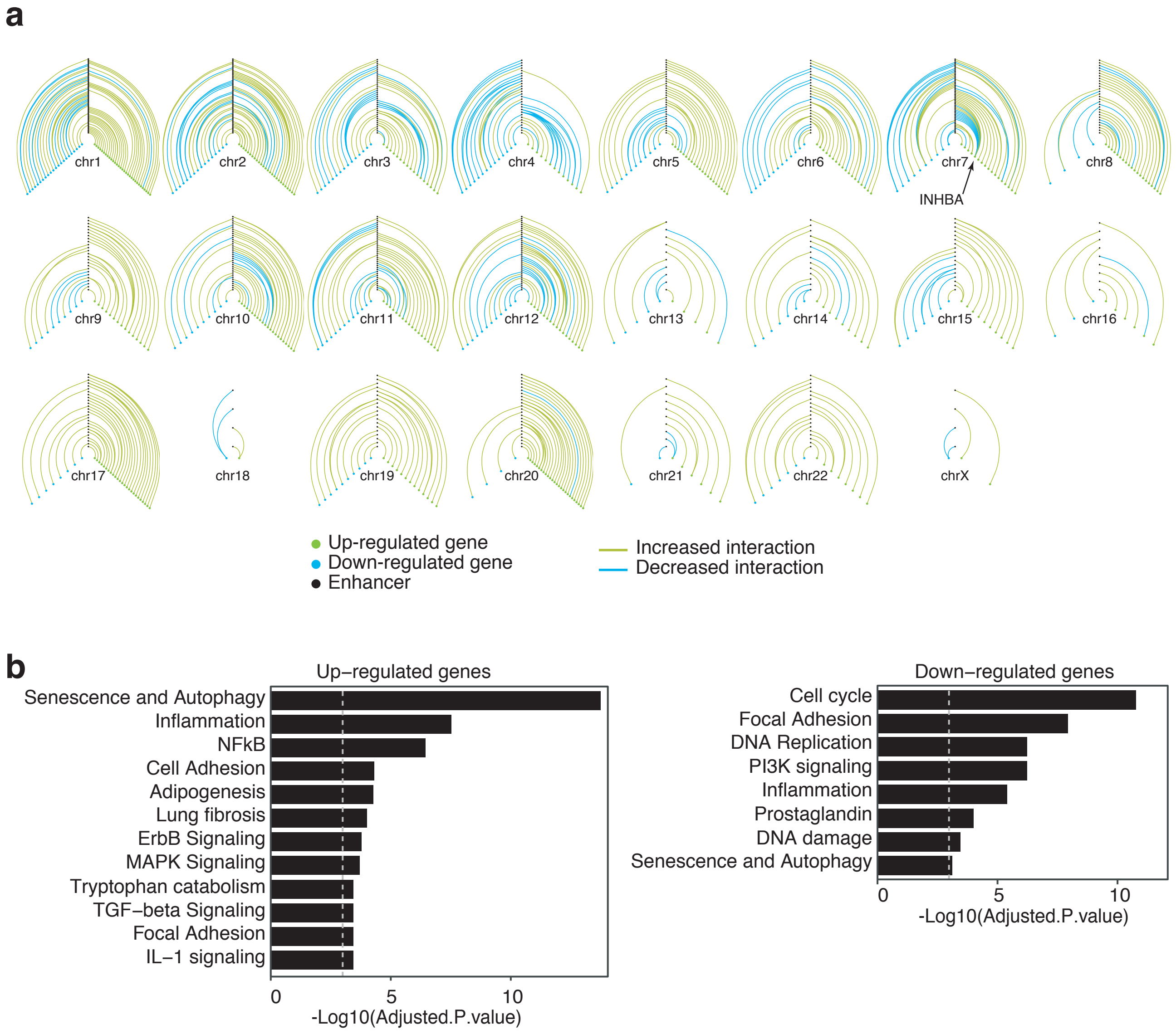


**Extended Data Figure 5. Genome-wide Hi-C EP differential network: a**, EP network using the filtering strategy based on EP changes annotated from cHi-C (Extended Data Fig. 4b, enhancers larger than 7.5 kb and resolution higher than 30 kb, i.e. bin sizes smaller than 30 kb) on each chromosome, with the vertical axis representing enhancers and the right and the left axes corresponding to up-regulated and down-regulated genes, respectively. **b**, Gene enrichment with EnrichR against the WikiPathways 2019 database of up-regulated (left) and down-regulated (right) genes in the genome-wide Hi-C EP differential network; the dotted grey line corresponds to 0.05 adjusted p-value, the selected threshold for significant enrichment. Terms were manually simplified and obvious redundancy was removed.

### Extended Data Fig. 6_Olan et al.

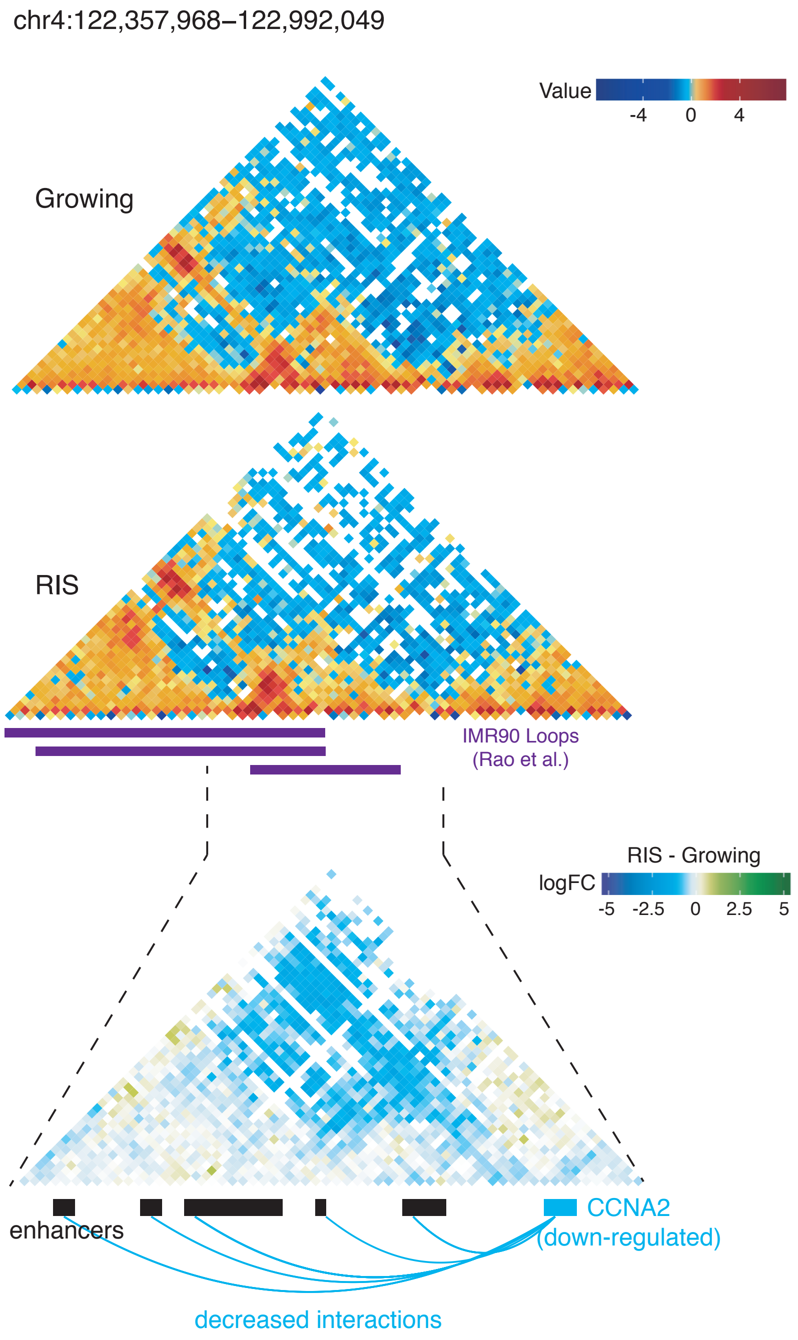


**Extended Data Figure 6. Down-regulation of CCNA2 associated with decreased EP interactions:** Growing and RIS Hi-C matrices (10 kb resolution) centred on the IMR90 loop (Rao et al.) consisting of the CCNA2 gene promoter and associated enhancers, as well as cHi-C differential log-fold change matrix (5 kb resolution) of this loop and significant decreased interactions (blue arcs) between the *CCNA2* gene promoter and associated enhancers.

### Extended Data Fig. 7_Olan et al.

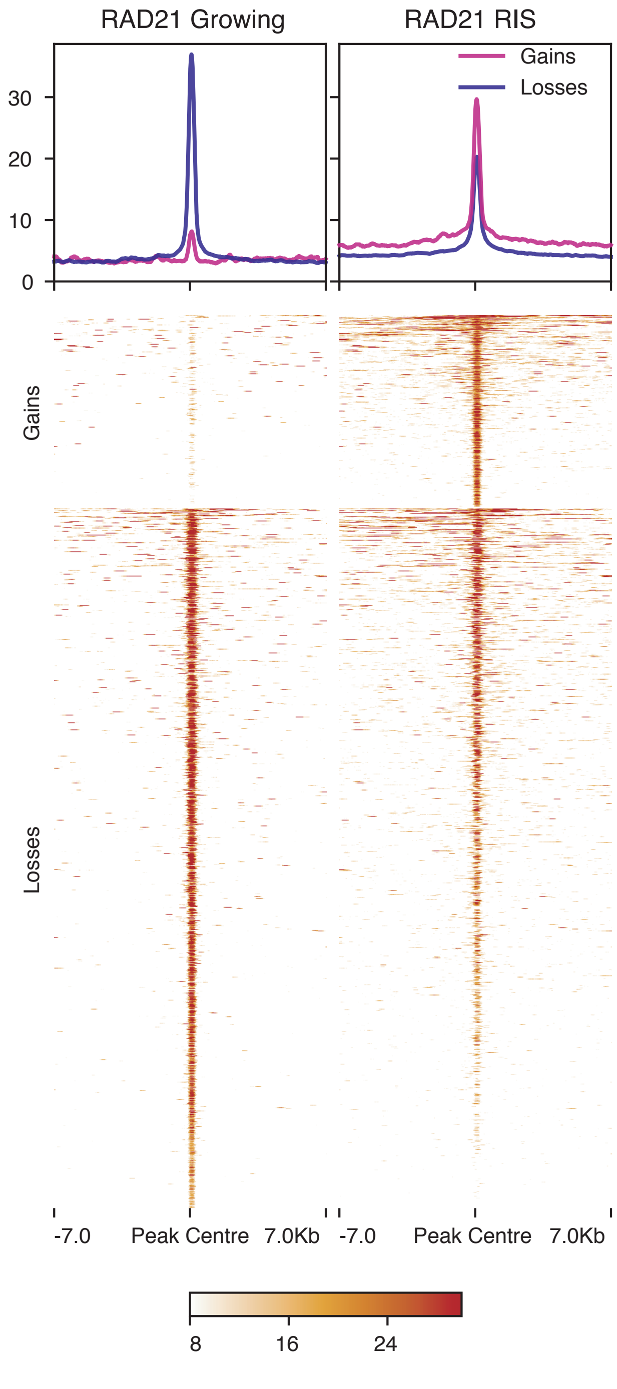


**Extended Data Figure 7. RAD21 binding gains and losses in growing and RIS:** Profiles and heatmaps of THOR-normalised RAD21 ChIP-seq signal in growing and RIS, centred on peaks overlapping regions with significant increased or decreased binding, as determined with THOR at FDR 0.05.

### Extended Data Fig. 8_Olan et al.

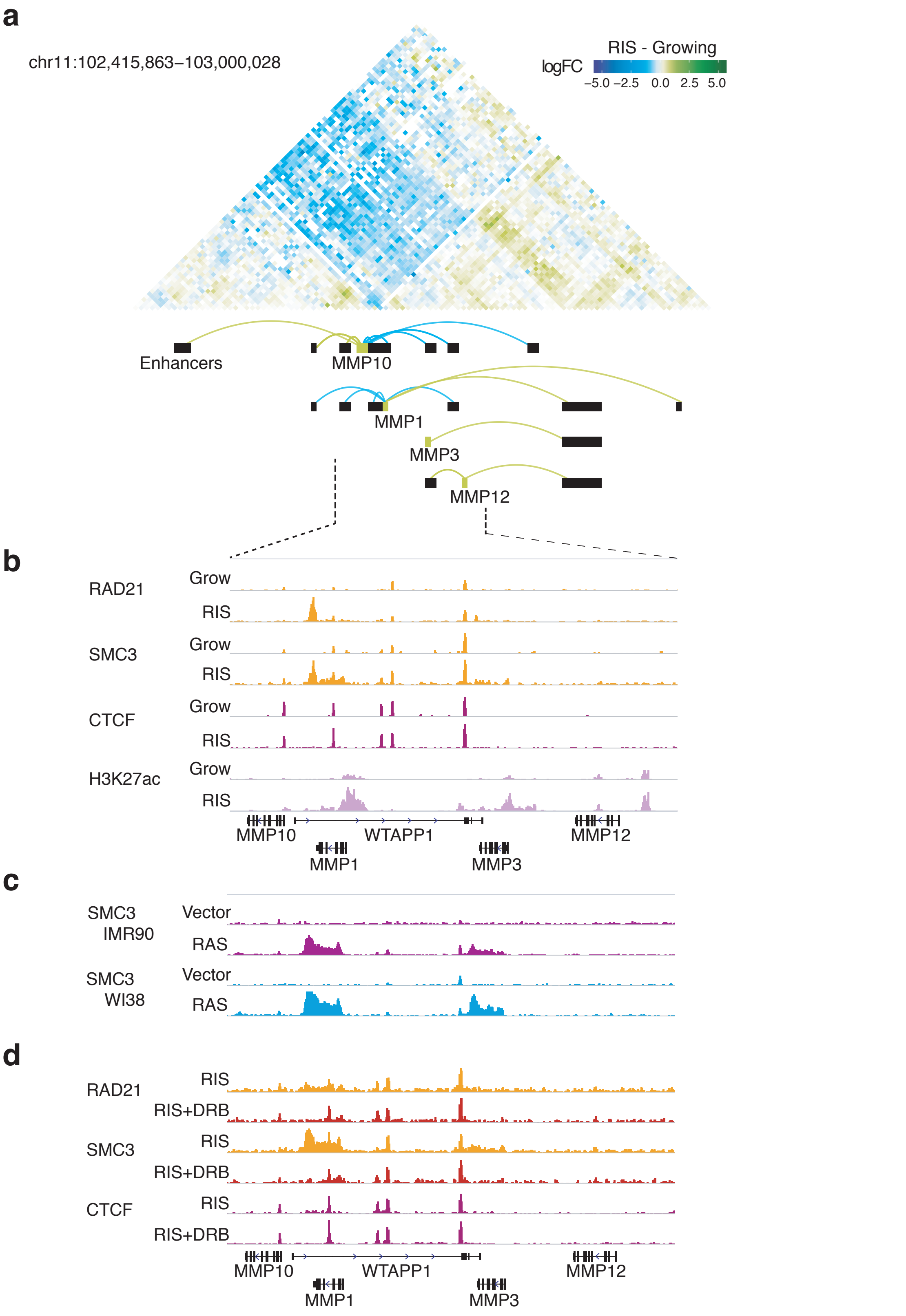


**Extended Data Figure 8: Interaction changes during RIS at the *MMP* locus suggesting increased spatial separation around *MMP1*: a**, cHi-C differential interaction matrix (HindIII resolution) at the *MMP* locus, consisting of the promoters of *MMP10*, *MMP1*, *MMP3*, *MMP12* and associated enhancers, as well as the significant EP contacts between them (green and blue arcs, corresponding to increased and decreased interactions, respectively). **b**, ChIP-seq THOR-normalized tracks at the *MMP* locus marked in **a**, with dotted lines of RAD21, SMC3, CTCF and H3K27ac in growing and RIS, as well as the *MMP* genes. **c**, SMC3 CHIP-seq of the *MMP* locus in RIS IMR90 and WI38 cells via constitutive expression of oncogenic HRAS-G12V and matched growing controls (empty vector). **d**, RAD21, SMC3 and CTCF ChIP-seq in RIS with and without DRB treatment (transcription elongation inhibitor) at the *MMP* locus with cohesin islands at the end of *MMP1* and *MMP3*.

### Extended Data Fig. 9_Olan et al.

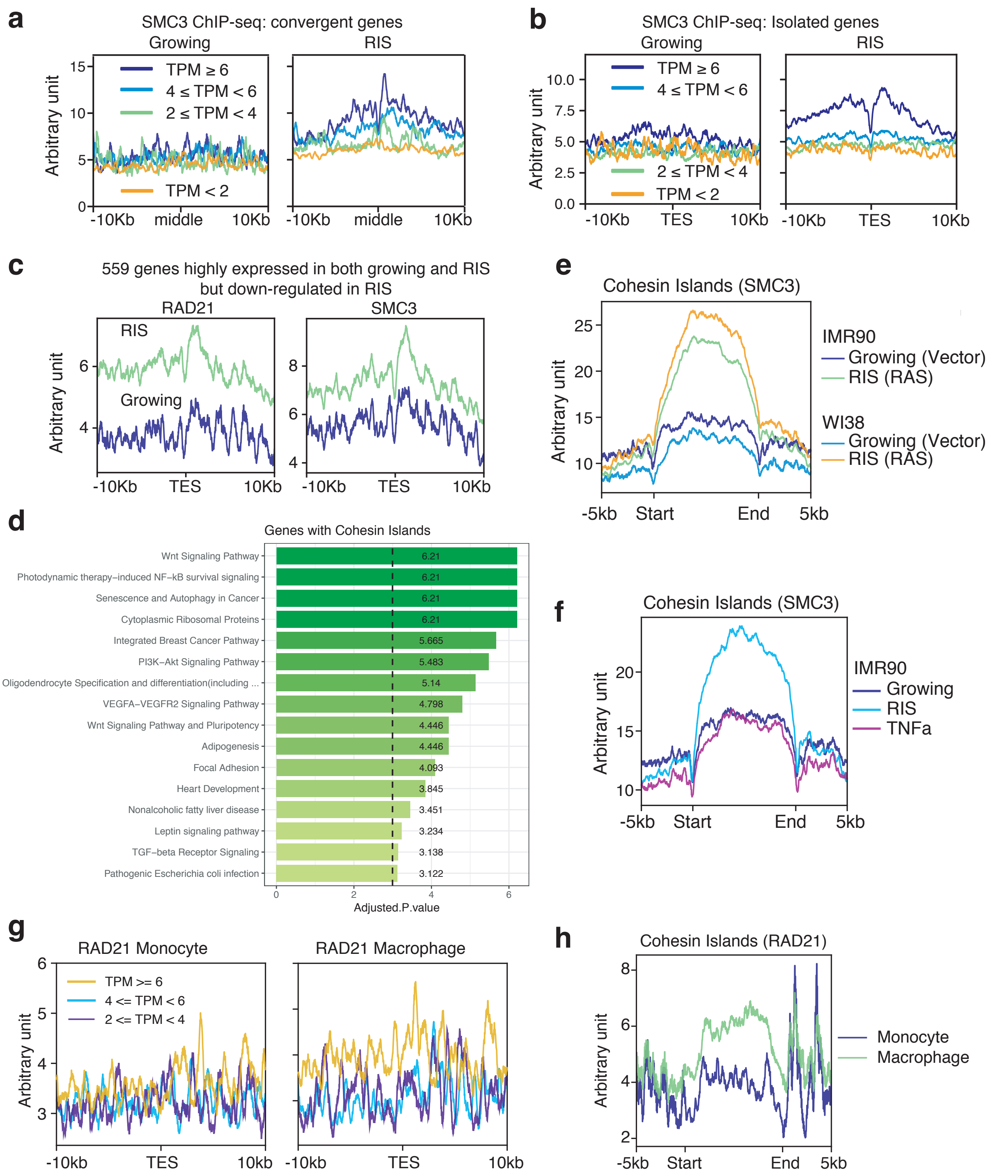


**Extended Data Figure 9: Cohesin islands in RIS IMR90 cells and THP-1 macrophages: a,** Distribution of SMC3 ChIP-seq signal between the 3' ends of convergent gene pairs, where both genes are in one of the four expression categories defined based on their log-transcripts-per-million (TPM* signifies TPM of both genes in the pair) in growing and RIS, respectively. **b**, SMC3 signal around 3' end (TES) of isolated genes averaged by four expression groups defined by log-TPM of the genes in growing and RIS, respectively. **c**, RAD21 and SMC3 ChIP-seq signal at the 3' end of 559 genes which are highly expressed in growing and RIS but more highly in growing. d, Gene enrichment pathways of genes with cohesin islands against WikiPathways 2019 (dotted line corresponds to the significance threshold, 0.05 p-adjusted). **e**, SMC3 ChIP-seq signal of constitutive HRAS-G12V-induced senescence and matched controls (empty vector), in IMR90 and WI38 cells, over the cohesin islands identified from RIS compared with RIS with DRB treatment. **f**, SMC3 ChIP-seq signal of RIS, TNFα-treated and matched control IMR90 cells over the cohesin islands. **g**, RAD21 ChIP-seq by Heinz et al. around the 3' end of genes grouped by expression levels (RNA-seq from Phanstiel et al.) in THP-1 monocytes and PMA-induced macrophages. **h**, Monocyte and macrophage RAD21 ChIP-seq by Heinz et al. over the cohesin islands defined in RIS IMR90 cells.
